# The SOCE system is critical for membrane bleb formation to drive avian primordial germ cell migration

**DOI:** 10.1101/2023.06.12.544577

**Authors:** Mizuki Morita, Manami Morimoto, Takayuki Teramoto, Junichi Ikenouchi, Yuji Atsuta, Daisuke Saito

## Abstract

Amoeboid cell migration is driven by the specialized cell protrusion, membrane bleb. A recent *in vitro* analysis of bleb formation using a cancer cell line showed that store-operated calcium entry (SOCE)-mediated local elevation of Ca^2+^ concentration triggers bleb formation, but it remains unknown how commonly this system is utilized in the bleb formation and bleb-driven cell migration *in vivo*. We demonstrate with avian primordial germ cell (PGC) model that that chick PGCs use SOCE, triggered by stem cell factor (SCF), to induce bleb formation and that it is essential for *in vivo* migration including trans-endothelial and mesenchymal cell migration. Our discovery also provides insight into the correlation between cancer metastasis/invasion and SOCE-mediated bleb formation.

## INTRODUCTION

Amoeboid cell movement is a mode of cell migration widely utilized in various cell types^1, 2^, including the protozoan ameba, normal motile cells such as the primordial germ cells and immune cells in vertebrates^3, 4^, and metastatic cancer cells^5, 6^. This universal mode of cell migration is driven by the formation of membrane blebs, which are balloon-like spherical protrusions of the plasma membrane (PM) at the front of migrating cells. Recent studies have revealed that the a local rise in calcium ion (Ca^2+^) concentration^3^, cortical actomyosin contraction^3^, plasma membrane detachment from the underling actin filaments^7, 8^, and influx of cytoplasm into blebs by the hydrostatic pressure^9, 10^, are critical for bleb formation. Importantly, bleb induction is thought to be initiated by a restricted rise of Ca^2+^ concentration to cause these other events^3, 9^. Recently, Aoki *et al.* showed that this earliest event during bleb induction is regulated via store-operated calcium entry (SOCE) in cancer cell lines *in vitro^9^*. However, it remained to be resolved whether all blebs are formed via SOCE.

Primordial germ cells (PGCs), which are germ cell precursors in the embryo, are formed in the extraembryonic or peripheral embryonic region and migrate long distance to the gonadal region of the embryo proper during animal development^11, 12, 13^. During their migration, PGCs exhibit bleb-type protrusion in a wide variety of animal embryos^3, 14, 15, 16, 17^, and a previous study using a zebrafish model demonstrated that blebs play a role in promoting their locomotion^3, 18^. In the context of zebrafish PGC migration, the local elevation of Ca^2+^ concentration is also critical to initiate bleb formation^3^. However, how the Ca^2+^ elevation occurs in PGC blebs remains unknown. In this study, we demonstrate with chick PGC model that SOCE is responsible for elevating Ca^2+^ concentration in forming blebs and SOCE-mediated bleb formation is essential for normal migration *in vivo*.

## RESULTS

### Elevation of Ca^2+^ concentration and disappearance of actomyosin cortex occur in the expanding blebs in chick PGCs

To characterize the bleb of chick PGCs, we first examine two signatures that define the bleb; dynamics of Ca^2+^ concentration and actomyosin cortex. It was reported that Ca^2+^ concentration rapidly rises in the expanding blebs^3, 9^, and actomyosin cortex disappears in the expanding blebs and is reformed in the retracting blebs during the bleb cycle^3, 8, 19, 20^. We have conducted the *in vitro* under-agarose assay, which enables to induce the blebs in cells by applying the pressure brought about by agarose weight (Fig. 1a)^21, 22^. This assay induces 2-5 large blebs per chick PGC (Fig. 1b). The cycle of each bleb is approximately 1min. To examine whether local Ca^2+^ elevation corresponding to bleb formation occurs within the blebs, we use GCaMP6s (Ca^2+^ sensor probe)-expressed PGCs in the under-agarose assay. This analysis demonstrates that a sharp rise in Ca^2+^ concentration occurs in the blebs during the expanding phase (Fig. 1c, d). In contrast, no change in Ca^2+^ concentration in the cell bodies is observed throughout the time course (Fig. 1d). Comparison of Ca^2+^ concentration between the blebs and cell body reveals that Ca^2+^ is significantly higher in the blebs than in the cell body (Fig. 1e). We also examine the dynamics of actomyosin cortex during the bleb cycle using PGCs with EGFP-tagged myosin regulatory light chain 1 (MRLC1) and Lifeact-mCherry (filamentous actin (F-actin) probe) by the *in vitro* under-agarose assay. Both MRLC1 and F-actin are not accumulated in the cortex of expanding blebs, and reappear during the retraction phase (Fig. 1f, g). These data indicate that chick PGCs also exhibit the typical dynamics of Ca^2+^ and actomyosin cortex during bleb formation.

**Fig. 1.**
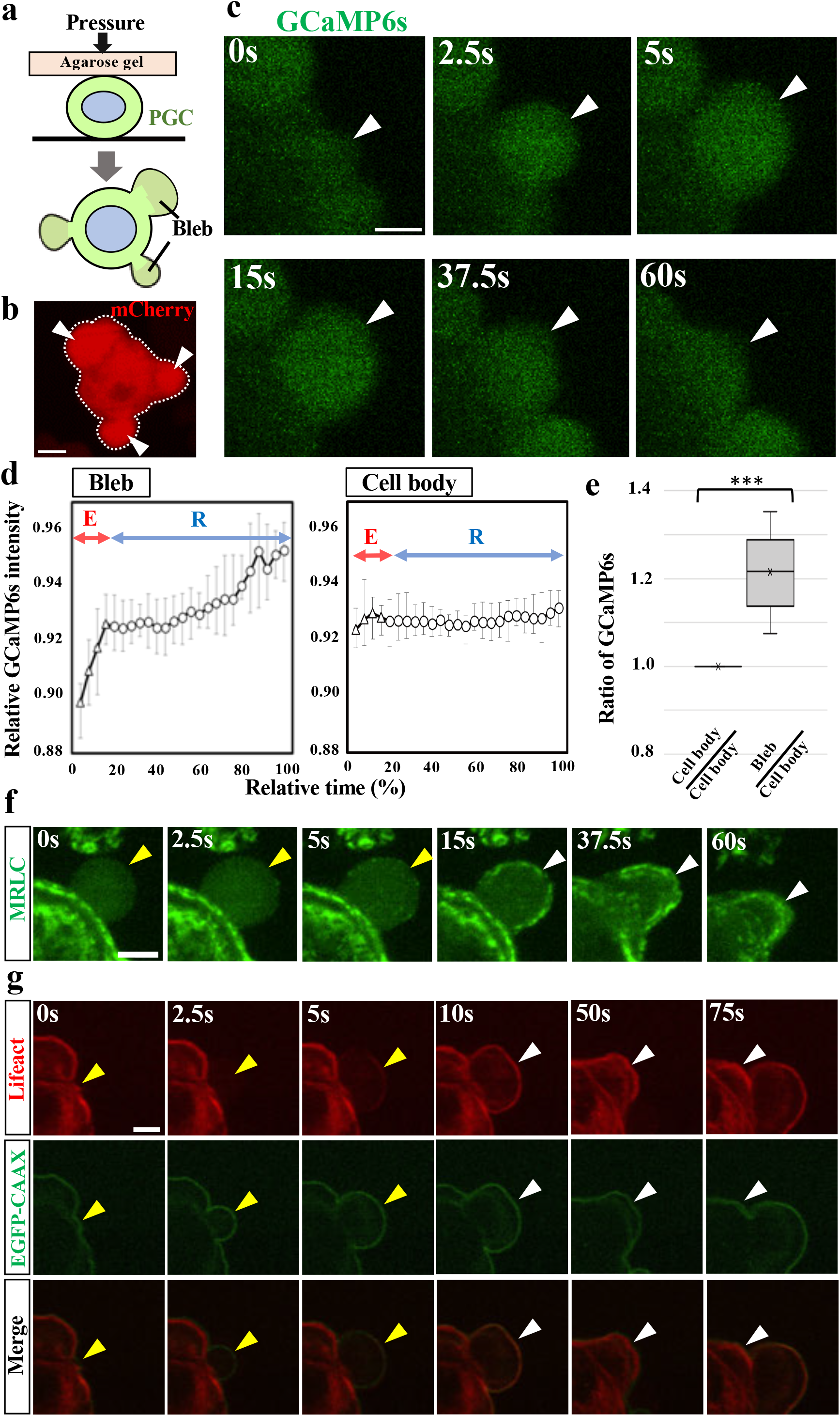
Dynamics of Ca^2+^ and actomyosin cortex in the expanding blebs of chicken PGCs. **a** The *in vitro* under-agarose assay applied to PGCs. To apply moderate pressure to PGCs, the polystyrene beads are placed with PGCs between the gel and dish. Several large blebs are induced. **b** A PGC in which the blebs (white arrowheads) are induced by this assay. mCherry is overexpressed. The cell outline is delineated by a white dotted line. **c** Motion captures from the movie (Supplementary Movie 1) corresponding to the bleb region (white arrowheads) in GCaMP6s (green)-expressing PGC during a typical bleb cycle. **d** Changes in GCaMP6s fluorescence intensities in the cytoplasm of the bleb regions (*N*=3) and cell bodies (*N*=3) during a bleb cycle. E and R indicate extending and retracting phases of bleb formation, respectively. Error bars indicate standard error. GCaMP6s fluorescence intensity in each region is corrected by fluorescence intensity of co-introduced cytoplasmic mCherry. **e** GCaMP6s fluorescence intensity ratio in the bleb and cell body under the *in vitro* under-agarose assay. *N*=10. ***p < 0.001 (Two-sided, unpaired Student’s *t* test). **f, g** Motion captures from the movie (Supplementary Movie 2, 3) corresponding to the bleb region in MRLC-EGFP (green in **f**)- or EGFP-CAAX + Lifeact-mCherry (green and red in **g**, respectively)-expressing PGC during a typical bleb cycle. Yellow and white arrowheads indicate the extending and retracting bleb, respectively. Scale bars: 2.5 μm in **b**, **c**, 3 μm in **f**.

### PGCs uptake extracellular Ca^2+^ via SOCE during bleb formation

We have sought to identify where is the source of Ca^2+^ to drive bleb formation. Cytosolic Ca^2+^ is mainly supplied from the extracellular environment and/or endoplasmic reticulum (ER)^23^. To verify whether extracellular Ca^2+^ is needed for bleb induction, we treat PGCs with a chelator of extracellular Ca^2+^, 1,2-bis (o-aminophenoxy) ethane N,N,N’,N’-tetraacetic acid (BAPTA) under the under-agarose assay. In this case, PGCs exhibit multiple tiny protrusions with low Ca^2+^ concentration instead of forming large blebs (Fig. 2a-c). Next, we investigate whether Ca^2+^ supply from the ER is involved in bleb formation treating PGCs with 2-aminoethyl diphenylborinate (2-APB), which is an IP3 receptor inhibitor resulting in blocking Ca^2+^ release from the ER to cytosol^24, 25^. Interestingly, this drug exerts a similar effect as BAPTA (Fig. 2a-c). These data indicate that influxes from both extracellular and ER are critical for bleb induction in chick PGCs. The Ca^2+^ uptake system using both influxes appears to be a feature of SOCE^26, 27^, which is the mechanism that takes Ca^2+^ from extracellular environment, triggered by Ca^2+^ depletion in the ER^28, 29^. When STIM proteins on the ER membrane sense the decrease of Ca^2+^ concentration, STIM proteins translocate to membrane contact sites between the ER and PM and then interact with and activate Orai1, a pore subunit of the calcium release-activated calcium channel (CRAC) on the PM to import Ca^2+^ from extracellular environment^30, 31^. Recent work with cancer cell line has reported that the ER contacts the PM in the blebs and activates Orai1 Ca^2+^ channels, resulting in elevation of Ca^2+^ concentration in the blebs^9^. To examine whether chick PGCs utilize SOCE to induce the blebs, we investigate the ER dynamics under the under-agarose assay using ER labeling dye. We have found that the tip of ER invades into the forming blebs in chick PGCs (Fig. 2d) as observed in the cancer cells^9^. Blocking SOCE by its inhibitor SKF96365 also causes multiple tiny protrusions with low Ca^2+^ concentration as shown in BAPTA and 2-APB treatments (Fig. 2a-c). Collectively, the under-agarose assay demonstrates that PGCs uptake extracellular Ca^2+^ via SOCE during bleb formation.

**Fig. 2.**
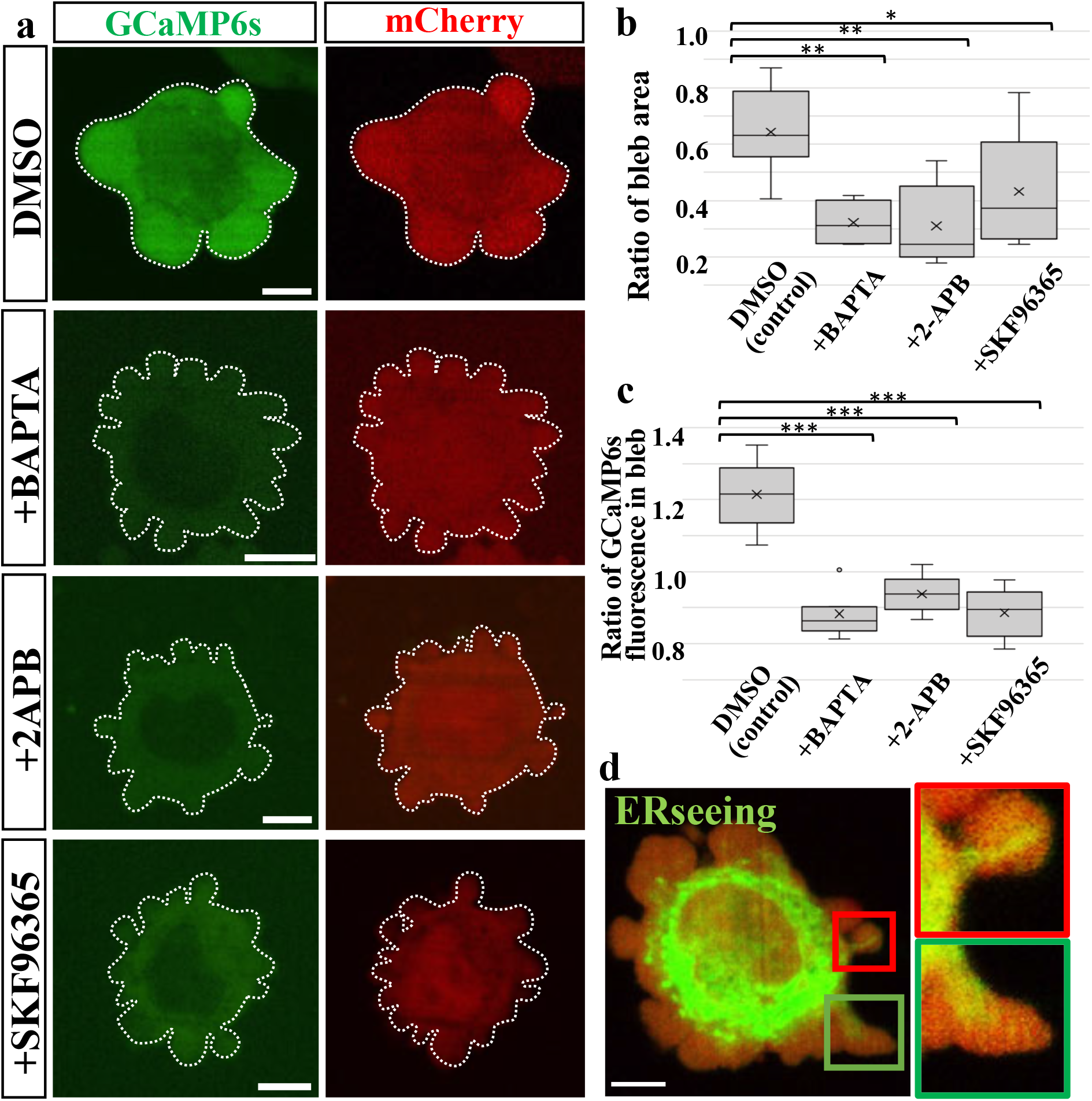
PGCs uptake extracellular Ca^2+^ by the SOCE system during bleb formation. **a** The pictures of GCaMP6s (green) and mCherry (red)-expressing PGCs when the *in vitro* under-agarose assay is performed in the presence of DMSO (control), BAPTA, 2-APB, and SKF96365. The cell outlines are delineated by a white dotted lines. **b, c** Ratio ofbleb area to total cell area **(b),** and ratio of GCaMP6s fluorescence intensity in the cytoplasm of the bleb to cell body **(c)** under the *in vitro* under-agarose assay. *N*=10 (DMSO), 7 (BAPTA), 5 (2-APB), 6 (SKF96365). GCaMP6s fluorescence intensity in each region is corrected by fluorescence intensity of co-introduced cytoplasmic mCherry. X indicates the mean value. **d** ERseeing staining (green) in Gap-mOrange (red)-expressing PGC. Rectangle regions enclosed by red and green lines are magnified in the right sides. *: P<0.05, **: P<0.01, ***: P<0.001 (Two-sided, unpaired Student’s *t* test). Scale bars: 5 μm in **a, d.**

### Stem cell factor promotes PGC motility *in vitro*

Next, we examine whether migrating PGCs exploit SOCE to induce the bleb and to drive locomotion. Since PGCs rarely form the blebs under the normal culture condition, we have attempting to develop the *in vitro* migration assay system for chick PGCs. We focus on two extracellular ligands, stromal derived factor-1 (SDF-1) and stem cell factor (SCF), to promote PGC movement *in vitro*. SDF-1 has been shown to guide PGC migration in zebrafish embryos^32^, and SCF has been shown to promote PGC motility in mouse embryos^33^. Chick PGCs express CXCR4 and c-Kit, their cognate receptors, respectively ^34, 35^.

Almost all PGCs rarely translocate when they are cultured alone or they are co-cultured with Cos7 cells expressing Gap-TdTomato or SDF-1 in Matrigel (Fig. 3a-d and Supplementary Fig. 1). In contrast, more than half of PGCs drastically migrate when co-cultured with Cos7 expressing SCF (Fig. 3a-e). In this co-culture set-up, Cos7 cells are placed on the right side relative to PGCs, but no directional bias is observed with respect to PGC migration (Fig. 3b), suggesting that SCF involved in promotion of PGC motility, rather than directional migration *in vitro*. Interestingly, PGCs move forward with single bleb-like large protrusion at the cell front and keep it during migration phase. In conclusion, we have established the *in vitro* migration assay system suitable for analyzing chick PGC migration.

**Fig. 3.**
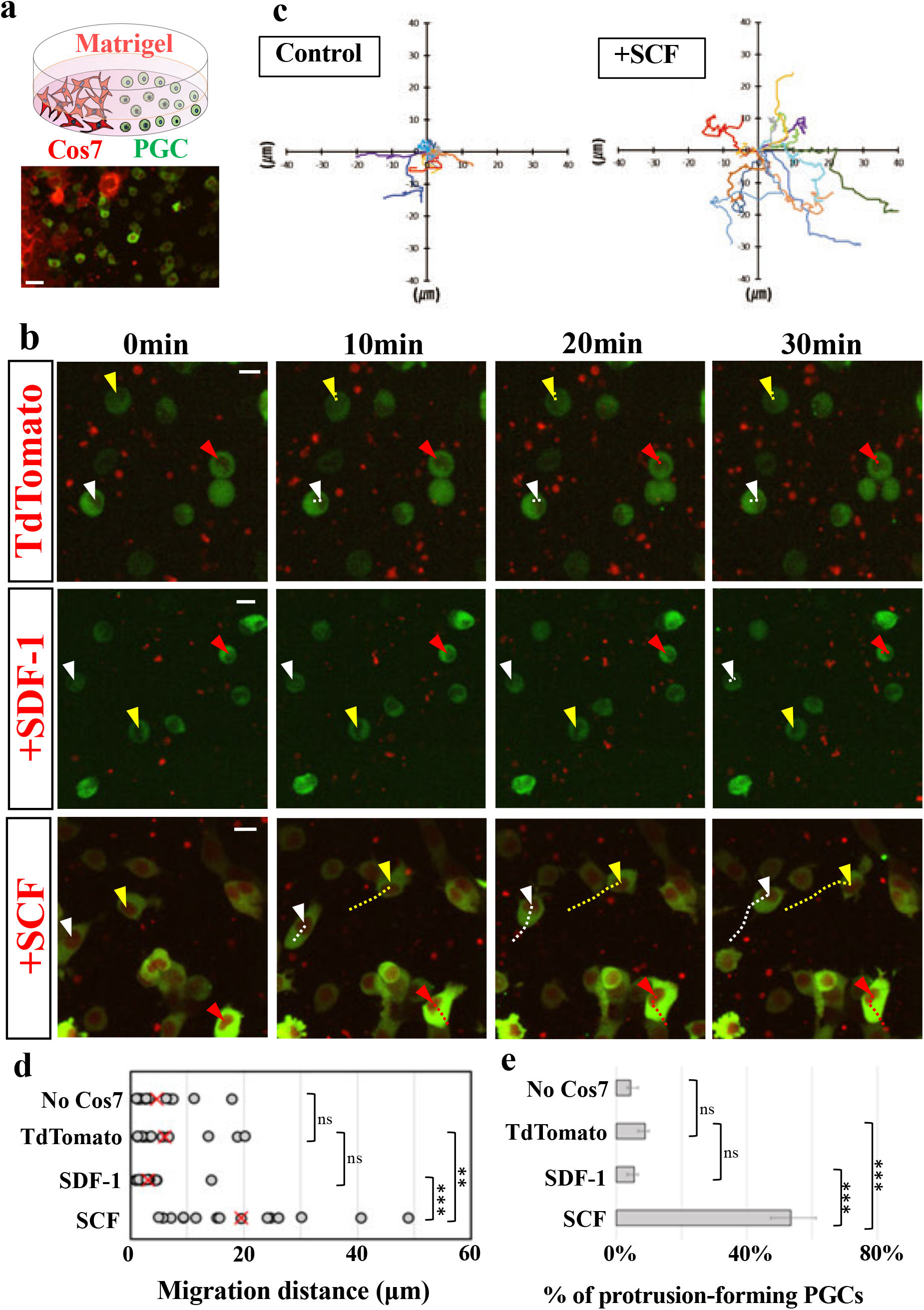
Stem cell factor promotes PGC motility *in vitro*. **a** Diagram and actual picture explaining co-culture of PGCs (GCaMP6s+, in green) and Cos7 cells (Gap-TdTomato+, in red) in Matrigel. **b** Motion captures from the movie (Supplementary Movie 4-6) taking GCaMP6s-expressing PGCs when co-cultured with Cos7 cells expressing GapTdTomato alone (control), SDF-1, and SCF, respectively. Arrowheads and dotted lines colored in each color indicate nuclei of randomly selected PGCs and their migration trajectories. **c** Summary of PGC migration trajectories when co-culture with Cos7 expressing Gap-TdTomato (control) alone or with Cos7 co-expressing SCF and Gap-TdTomato. Cos7 cells located in the left side. *N*=15 cells. **d, e** Migration distances of PGCs (*N*=15 cells) for 30 min and ratio of protrusion-forming PGCs (*N*=30) when co-cultured with no Cos7 cells, Gap-TdTomato-, SDF-1-, or SCF-overexpressing Cos7 cells. X and error bars indicate the mean value and standard error, respectively. **: P<0.01, ***: P<0.001 (Two-sided, unpaired Student’s *t* test). Scale bars: 30 μm in **a**, 15 μm in **b**.

### SOCE-driven bleb formation is essential for PGC migration

To investigate whether the large protrusions formed at the cell front have the characteristics of bleb, we observe actin, Ca^2+^ and the ER dynamics in PGCs in this *in vitro* migration assay under the condition with SCF. Lifeact-mCherry-expressing PGCs show that their large protrusions at the cell front are actin-free, while their cell bodies and rear regions have cortical F-actin and stress fiber-like actin bundles (Fig. 4a). The absence of actin in the cell front continues more than 30 min in some PGCs. In zebrafish model, the thick actin fiber structure at the base of the bleb, called actin brushes, is observed during bleb cycle^36^, but no actin brushes-like structures have been observed in chick PGCs. High GCaMP6s signal are also detected at the cell front compared to the rear end and continues throughout migration phase (Fig. 4b-d). ER staining exhibits invasion of the ER into the migration front (Fig. 4e). Collectively, these data indicate that PGCs in the assay form the bleb during migration phase.

**Fig. 4.**
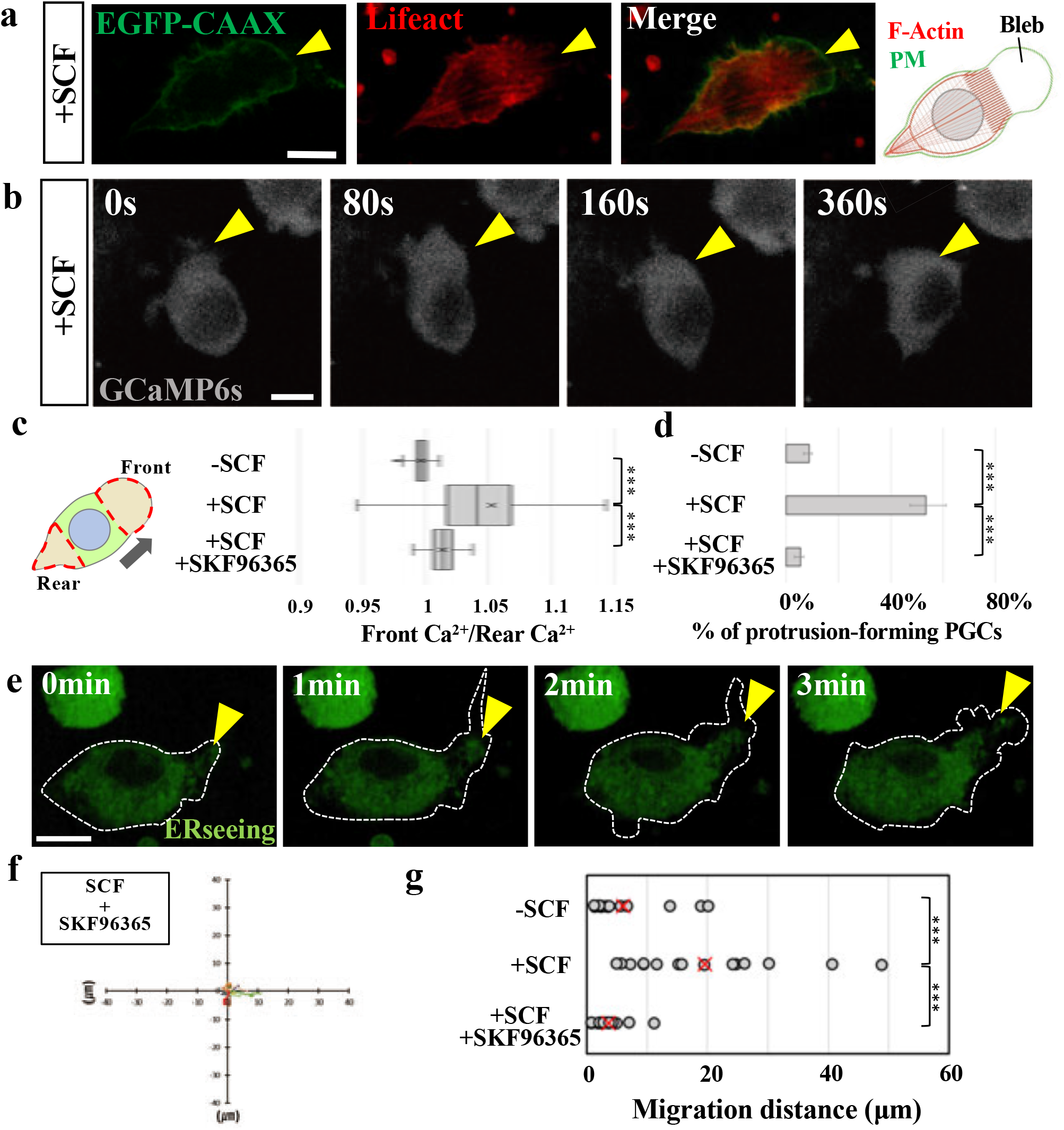
The SOCE system-driven bleb formation is essential for PGC migration. **a** The distribution pattern of EGFP-CAAX (green) and Lifeact-mCherry (red) in a migrating PGC under the *in vitro* migration assay with SCF. PM, plasma membrane, **b** Motion captures from the movie (Supplementary Movie 7) about a GCaMP6s-expressing migrating PGC under the condition with SCF. **c, d** Ratio of Ca^2+^ concentration (GCaMP6s fluorescence) at the front and rear end in migrating PGCs (*N*=10 cells. 20 measurements per cell) and ratio of protrusion-forming PGCs (*N*=30 cells) under the condition with/without SCF and SKF96365. Error bars indicate the standard error. **e** Motion captures from the movie (Supplementary Movie 8) taking a PGC stained by an ER tracker, ERseeing (green). The cell outline is delineated by a white dotted line. **f** Summary of PGC migration trajectory in the condition with Cos7 expressing SCF and SKF96365. *N*=15 cells. **g** Migration distances of PGCs for 30 min under the condition with/without SCF and SKF96365. *N*=15. Yellow arrowhead indicates the bleb formed at the cell front. ***: P<0.001 (Two-sided, unpaired Student’s *t* test). Scale bars: 10 μm in **a, b, e.**

Next, to verify whether PGC migration is dependent on SOCE, we treat PGCs in the migration assay with a SOCE inhibitor, SKF96365. As results, Ca^2+^ level at the cell front becomes low and its difference between the front and rear is reduced (Fig. 4d), no PGCs with the protrusion are observed (Fig. 4e), and PGC migration is drastically suppressed (Fig. 4f). Thus, SOCE-driven bleb formation is essential for chick PGC migration.

### The dominant negative mutant of Orai1 inhibits SOCE in PGCs in cell-autonomous manner

Prior to investigating the SOCE function *in vivo*, we prepare PGCs in which SOCE is impaired in a cell-autonomous manner. The dominant negative mutant of Orai1 (Orai1 E106Q) is Tet-on inducibly expressed in PGCs with Dox administration. These PGCs exhibit no blebs as with the addition of SKF96365 under the under-agarose assay (Fig. 5a, b). Orai1 E106Q-expressed PGCs show markedly lower motility than the control under the *in vitro* migration assay with SCF (Fig. 5c, d). Collectively, this genetic tool shows an effective inhibitory effect on SOCE.

**Fig. 5.**
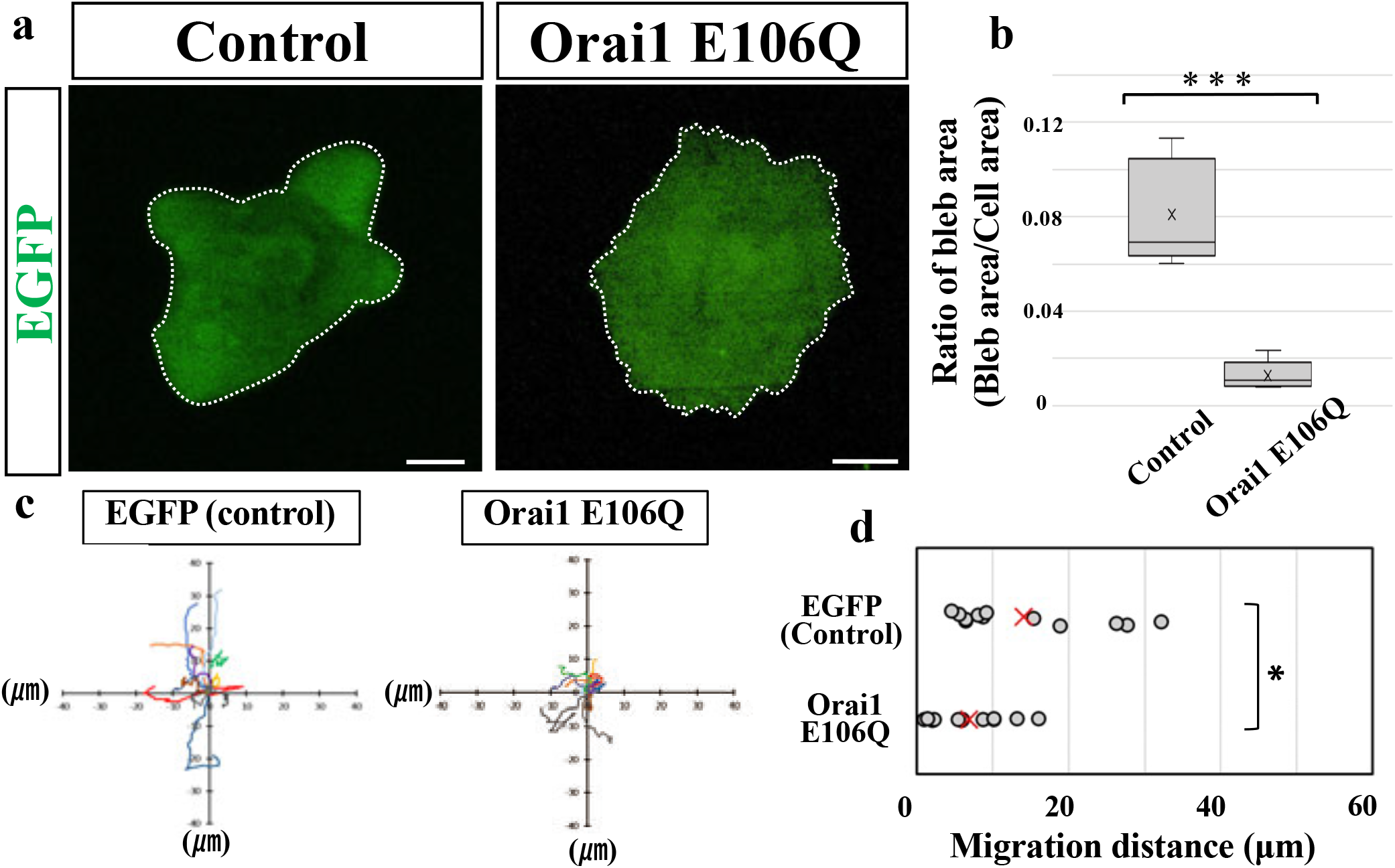
The dominant negative mutant of Orail inhibits the SOCE system in PGCs in cell-autonomous manner. **a** The pictures of EGFP (green) and Orai1 E106Q-overexpressing PGCs under the *in vitro* under-agarose assay. The cell outline is delineated by a white dotted line. **b** Ratio of bleb area to total cell area under the *in vitro* under-agarose assay. *N*=5 cells. **c, d** Summary of migration trajectories (*N*=12 cells) and migration distances for 30 min (*N*=12 cells) about PGCs expressing EGFP or Orai1 E106Q with EGFP under the *in vitro* migration assay with SCF. *: P<0.05, P<0.001 (Two-sided, unpaired Student’s *t* test). Scale bars: 5 μm in **a.**

### SOCE is also critical for PGC migration *in vivo*

Unlike mammals and teleost fish in which PGCs migrate within mesenchymal or epithelial tissue to reach the somatic gonadal primordia, PGCs in avian embryos exploit blood circulation to home to the gonadal primordia^37^. Thus, to examine whether chick PGCs exploit SOCE to migrate in the embryos, Orai1 E106Q+ (also EGFP+) PGCs with mCherry+ control PGCs (each 2,500 cells) are back-infused into the vascular system of host Hamburger and Hamilton (HH) stage 15 embryos. As a first step to reach the goal, circulating PGCs are normally arrested at a specific vascular plexus (extravasation vascular plexus; Ex-VaP) due to actin-dependent stiffness^38^. As a result, Orai1 E106Q+ PGCs are arrested in Ex-VaP as much as control PGCs 5 hrs after infusion (Fig. 6a, b), suggesting that SOCE is not responsible for PGC arrest.

**Fig. 6.**
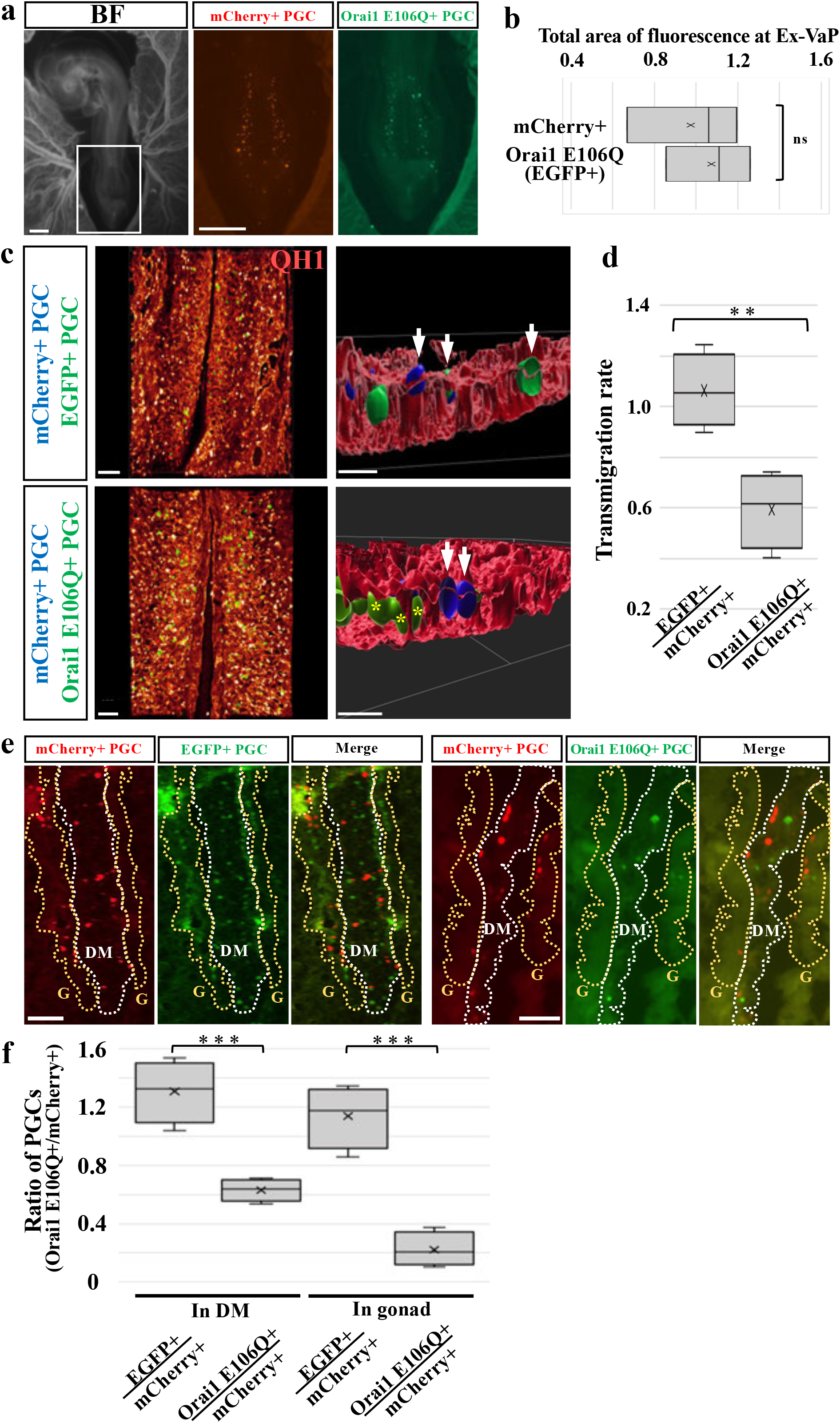
The SOCE system is also critical for PGC migration *in vivo*. **a** The distribution patterns of mCherry+ and Orai1 E106Q+ (also EGFP+) PGCs at HH 16 chick embryo after 5 hrs after infusion. Each is back-infused with 2,500 PGCs. Fluorescent images correspond to a square (Ex-VaP region) in the BF (bright field) image. **b** Quantification of arrested PGCs in Ex-VaP 5 hrs after infusion with 2,500 PGCs each. The total area occupied by each PGC line is calculated. **c** 3D reconstitution image of mCherry+ PGCs (blue) and EGFP (green) of Ex-VaP in HH16 quail embryo 6hrs after infusion. The vascular network of the embryo is immunostained by QH1 (red) and 3D-reconstracted. Left and right images are ventral and transverse view of Ex-VaP region, respectively. White arrows indicate transmigrating PGCs. White asterisks indicate remaining PGCs in intravascular space. **d** Ratio of transmigration rate of Orai1 E106Q+ PGCs. The number of EGFP+ (control) PGCs and Orai1 E106Q+ PGCs during and after transmigration is divided by the number of co-infused mCherry+ (infusion control) PGCs during and after transmigration, respectively. *N*=4 embryos. **e** The horizontal sections of E4.5 chicken embryos (HH25) 2 days after infusion with EGFP+ PGCs (green), Orai1 E106Q+ PGCs (green), and co-infused mCherry+ PGCs (red). White and yellow dotted lines delineate the dorsal mesenteric regions (DM) and gonadal primordium (G), respectively. **f** Ratio of the number of EGFP+ and Orai1 E106Q+ PGCs located in the DM and G 2 days after infusion. It is corrected by the number of co-infused mCherry+ PGCs. *N*=4 embryos. X and error bars indicate the mean value and standard error, respectively. **: P<0.01, ***: P<0.001 (Two-sided, unpaired Student’s *t* test). Scale bars: 1000 μm in **a**, 100 μm and 20 μm in left and right images of **c**, respectively, 100 μm in **e**.

Next, arrested PGCs transmigrate through an endothelial lining to the mesenteric mesenchyme during normal development. Previous reports demonstrated that PGCs insert their bleb-like protrusion into the vascular barrier and pass through it during transmigration phase^38, 39^. To examine whether SOCE is involved in this migratory process, we have conducted infusion of chick PGCs into HH15 quail embryos and immunostaining with QH1 (endothelial cell marker for quail) 6hrs after infusion (Fig. 6c, d). Quail embryos are used for this experiment with QH1 staining which is available only with this species. Indeed, the number of Orai1 E106Q+ (also EGFP+) PGCs during and after transmigration through an endothelial lining of Ex-VaP is reduced by 40% compared to control mCherry+ PGCs 6hrs after infusion (Fig. 6c, d). In case of co-infusion of EGFP+ and mCherry+ PGCs as an infusion control experiment, transmigration rates between these two cell lines are nearly equal (Fig. 6c, d). Thus, these data suggest that the disfunction of SOCE in PGCs leads to transmigration failure at Ex-VaP.

After transmigration, PGCs normally migrate through the dorsal mesentery to the gonadal primordium at around E4.5. After 2-day post-infusion stage corresponding to E4.5, a considerable number of control PGCs (EGFP+ and mCherry+) is found in the dorsal mesenteric and gonadal regions (Fig. 6e, f). In contrast, there is fewer Orai1 E106Q+ PGCs in these regions than controls (Fig. 6e, f). Interestingly, ratio of Orai1 E106Q+ PGCs to mCherry+ PGCs in the gonad (0.22) is lower than that in the dorsal mesentery (0.63). SOCE inhibition does not appear to suppress cell survival and proliferation of PGCs *in vitro* (Supplementary Fig. 2). These data suggest that SOCE is also necessary for PGC migration through the dorsal mesentery to the gonadal primordium rather than for cell survival and growth during this migration phase.

## DISCUSSION

Although reports that SOCE is required for cancer invasion^40^ and metastasis^41^ suggested that it is necessary for cell migration, it was unclear what role SOCE plays in cell migration. Recently, Aoki *et al.* revealed that SOCE is a central mechanism to regulate bleb formation in cancer cell line *in vitro*^9^. The remaining important issue is whether SOCE regulates bleb formation to drive cell migration *in vivo*. We first demonstrate that SOCE is a critical mechanism to regulate bleb formation to migrate *in vivo*.

The present study suggested that SOCE is required for both the PGC transmigration from the intravascular to the extravascular dorsal mesenteric zone and their migration through mesenchyme of the dorsal mesentery to the gonads (Fig. 6). Since PGCs form bleb-like protrusion during transmigration in the chick model^38^ and migration in the dorsal mesentery in the mouse model^42^, it is reasonable that SOCE would regulate the bleb formation to drive locomotion in these *in vivo* environments. We also demonstrate that SCF induces bleb formation via SOCE resulting in promotion of PGC motility *in vitro* (Fig. 3 and 4). SCF is expressed in the dorsal mesentery^43^, and promote the motility of mouse PGCs *in vivo*^42^. Previous works reported that this ligand induces reorganization of actin cytoskeleton^44, 45^. Collectively, SCF would induce the bleb formation in PGCs in the *in vivo* environments, and the remaining important issue is how SCF regulates SOCE.

We demonstrate that bleb formation is regulated via SOCE in avian PGCs. Previous study using the zebrafish model suggests that the elevation of Ca^2+^ concentration is necessary and sufficient to form bleb^3^, consistent with this study using the chicken model. However, the mechanism of the elevation of Ca^2+^ concentration in the bleb may differ between the two animal models. In zebrafish PGCs, the ER is confined to the rear side of the migrating PGCs and is not found within the bleb forming at the cell front^36^, suggesting that SOCE is not utilized to form bleb. In zebrafish PGCs, thick actin fiber structure, called actin brushes, is formed as if separating the cell body from the bleb of the cell front, raising the possibility that the actin brushes block the invasion of ER into the bleb^36^. Indeed, no actin brushes-like structures have been observed at the bleb base in migrating chick PGCs. Distinct mechanisms to raise Ca^2+^ concentration in the bleb between two species would result from differential regulations of actin cytoskeleton in these species. This study raises an interesting question, how the mechanism underling bleb formation in PGCs has been evolved and diversified among animal species.

The unanswered important questions are whether SOCE triggers or promotes bleb formation, how the bleb forming region is determined/restricted, and how the bleb persists during migration. We have established an effective analytical system to address these issues.

## MATERIALS and METHODS

### Animals, staging, and animal care

Fertilized chicken (*Gallus gallus domesticus*, White leghorn) eggs and fertilized quail (*Coturnix japonica*) eggs were purchased from Yamagishi poultry farm (Mie, Japan) and from Nagoya University through the National Bio-Resource Project of the MEXT, Japan, respectively. Eggs were incubated at 38.5℃, and embryos were staged by Hamburger and Hamilton’s stage^46^. All animal experiments were performed with the approval of the Institutional Animal Care and Use Committees at Kyushu University.

### Plasmid constructions

pT2A-BI-Gap-TdTomato-TRE: pT2AL200R150G vector^47^ was digested with XhoI-BglII. These sites were blunt-ended, and inserted with the fragment of pBI-TRE (Clontech) containing the bidirectional tetracycline-responsive element (TRE) with two minimal promoters of CMV in both directions, and two polyA-additional sequences of the rabbit beta globin gene. This vector was designated as pT2A-BI-TRE. The full-length of Gap-TdTomato^48^ was amplified by PCR, and subcloned into the SpeI-HindIII site of pT2A-BI-TRE. pT2A-BI-Gap-TdTomato-TRE- (SCF or SDF-1): The ORF of chicken SCF or chicken SDF-1^49^ was subcloned into the MluI-EcoRV site of pT2A-BI-Gap-TdTomato-TRE. The ORF of SCF was amplified and isolated from cDNA derived from E2.5 whole chick embryo by PCR. pT2A-BItight-EGFP-TRE: pT2AL200R150G vector was digested with XhoI-BglII. These sites were blunt-ended, and inserted with the fragment of pBItight-TRE (Clontech) containing the improved bidirectional tetracycline-responsive element (TRE) with two minimal promoters of CMV in both directions, and two polyA-additional sequences of the rabbit beta globin gene. This vector was designated as pT2A-BItight-TRE. The full-length of EGFP was amplified by PCR, and subcloned into the EcoRI-PstI site of pT2A-BItight-TRE. pT2A-BItight-EGFP-TRE-Orai1 E106Q: The ORF of human Orai1 with E106Q mutation and myc tag (addgene #22754) was amplified by PCR, and subcloned into the MluI-EcoRV site of pT2A-BItight-EGFP-TRE. pT2A-BItight-mCherry-TRE: The full-length of mCherry was amplified by PCR, and subcloned into the EcoRI-BglII site of pT2A-BItight-TRE. pT2A-BItight-GCaMP6s-2AP-mCherry: The consecutive sequences of GCaMP6s-2AP-mCherry^50^ was amplified by PCR, and subcloned into the EcoRI site of pT2A-BItight-TRE by In-Fusion^®^ HD cloning kit (TaKaRa). pT2A-BItight-Lifeact-mCherry-TRE: Lifeact-mCherry was amplified by PCR, and subcloned into the MluI-EcoRV site of pT2A-BItight-TRE. pT2A-CAGGS-Gap-mOrange: The sequence of Gap-mOrange^51^ was amplified by PCR, and subcloned into the MluI-EcoRV site into pT2A-CAGGS^38^. pT2A-CAGGS-EGFP-CAAX-IRES2-PuroR: The sequences of IRES2 and PuroR were amplified by PCR, and subcloned into the XhoI-EcoRI site and EcoRI-BglII site into pT2A-CAGGS, respectively. This vector was designated as pT2A-CAGGS-IRES2-PuroR. The full-length of EGFP-CAAX was amplified by PCR, and subcloned into the PstI-XhoI site of pT2A-CAGGS-IRES2-PuroR. pT2A-CAGGS-MRLC1-EGFP-IRES-PuroR: MRLC1-EGFP was amplified by PCR, and subcloned into the NotI-XhoI site of pT2A-CAGGS-IRES2-PuroR. pT2A-CAGGS-Tet3G-IRES2-PuroR: The full-length of Tet3G was amplified by PCR, and subcloned into the NotI-XhoI site of pT2A-CAGGS-IRES2-PuroR.

Primer sequences are as described below.

Gap43-SpeI-F: agaactagtggatccccgcggATGTTGTGCTGTATGAGAAGAACCAAGC

TdTomato-HindIII-R: gacaagcttgaattcTTACTTGTACAGCTCGTCCATGCCGTACAG

SDF-1-MluI-F: cagacgcgtATGGACCTCCGCGCCCTGGCTCTGCTCGCCT

SDF-1-EcoRV-R: agagatatcTTACTTGTTTAAAGCTTTCTCCAGATATTCC

SCF-MluI-F: aattacgcgtATGAAGAAGGCACAAACTTGGATTATCAC

SCF-EcoRV-R: aattgatatCTACACTTGTAGATGTTCTTTTTCTTTTTGCTGCAACA

EGFP-EcoRI-F: ggagaattcATGGTGAGCAAGGGCGAGGAGCTGTTCACCG

EGFP-PstI-R: agactgcagTTACTTGTACAGCTCGTCCATGCCGAGAGT

Orai1 E106Q-MluI-F: ctgacgcgtACCATGCATCCGGAGCCCGCCCCGCCCC

Orai1 E106Q-EcoRV-R: ggagatatcCTAGAGATCTTCCTCAGAAATGAGC

mCherry-EcoRI-F: ggagaattcATGGTGAGCAAGGGCGAGGAGGATAACATGG

mCherry-BglII-R: caggagatctTTACTTGTACAGCTCGTCCATGCCGCCGGT

GCaMP6s-inf-F: gtcagatcgcctggaATGGGTTCTCATCATCATCATCATCATGGTATGG

mCherry-inf-R: gactgcagcctcaggagatcTTACTTGTACAGCTCGTCCATGCCGCCGG

Gap43-MluI-F: attacgcgtcggcaccATGCTGTGCTGTATGAGAAGAACCAAGCAGGTG

mOrange-EcoRV-R: tctgataTCACTTGTACAGCTCGTCCATGCCGCCG

IRES2-XhoI-F: aatctcgagGCCCCTCTCCCTCCCCCCCCCCTAAC

IRES2-EcoRI-R: aatgaattcTGTGGCCATATTATCATCGTGTTTTTCA

PuroR-EcoRI-F: agagaattcATGACCGAGTACAAGCCCACGGTGCGCCTCG

PuroR-BglII-R: aaaagatctTCAGGCACCGGGCTTGCGGGTCATGCACCAGGT

EGFP-PstI-F: aagctgcagcggcaccATGGTGAGCAAGGGCGAGGAGCTGTTCACCGGG

CAAX-XhoI-R: gcctcgagTTACATAATTACACACTTTGTCTTTGACTTCTTTTTCTTCT

MRLC1-NotI-F: tgcagcggccgcATGTCCAGCAAGCGGGCCAA

EGFP-XhoI-R: gggcctcgagTTACTTGTACAGCTCGTCCATG

Tet3G-NotI-F: gcagcggccgcATGTCTAGACTGGACAAGAGCAAAGTC

Tet3G-XhoI-R: ggcctcgagTTACCCGGGGAGCATGTCAAGGTCAAAAT

### Establishment and maintenance of PGCs in culture

Circulating PGCs along with blood cells were harvested from blood of HH 15 chicken embryos, and were cultured in calcium-free DMEM (Gibco) diluted with water, containing <u>F</u>GF2 (Wako), <u>A</u>ctivin A (APRO Science), and <u>c</u>hicken <u>s</u>erum (FAcs medium) according to the method previously described ^52, 53^. After one month, expanded PGCs were cryo-preserved at -80 ℃ in Bambanker (NIPPON Genetics) until used for experiments.

### Plasmid transfection and establishment of gene-manipulated PGC lines and Cos7

5 x 10^4^ cultured PGCs were washed with OPTI-MEM (Gibco) and placed in 96 well plate containing 100 μl of FAot medium^53^ (adding Ovo-transferrin instead of chicken serum) without heparin and antibiotics (penicillin, streptomycin, and amphotericin) for 3hrs. Transfection mix containing 0.2 μg of total plasmids and 0.5 μl of Lipofectamine 2000 (ThermoFisher Sciencetific) in 50 μl of OPTI-MEM were added to PGCs in 96 well plate. The medium was replaced by conventional FAcs medium containing heparin and antibiotics 6hrs after addition of transfection mix. We used 7 different sets of plasmids (pT2A-BItight-GCaMP6s-2AP-mCherry + pT2A-CAGGS-Tet3G-IRES2-PuroR + pCAGGS-T2TP^54^; pT2A-CAGGS-Gap-mOrange + pT2A-CAGGS-MRLC-EGFP-IRES-PuroR + pCAGGS-T2TP; pT2A-BItight-Lifeact-mCherry-TRE + pT2A-CAGGS-Tet3G-2AP-PuroR^38^ + pT2A-CAGGS-EGFP-CAAX-IRES2-PuroR + pCAGGS-T2TP; pT2A-CAGGS-Gap-mOrange + pCAGGS-T2TP; pT2A-BItight-EGFP-TRE + pT2A-CAGGS-Tet3G-IRES2-PuroR + pCAGGS-T2TP; pT2A-BItight-mCherry-TRE + pT2A-CAGGS-Tet3G-IRES2-PuroR + pCAGGS-T2TP; pT2A-BItight-EGFP-TRE-Orai1 E106Q + pT2A-CAGGS-Tet3G-IRES2-PuroR + pCAGGS-T2TP) for transfection to PGCs. Following transfection, PGCs were cultured in the FAcs medium containing 0.1-0.3 μg/ml puromycin for 3 days to enrich puromycin-resistant cells.

Cos7 cells were maintained at 37℃ with Dulbecco’s modified Eagle’s medium (DMEM) containing 1.5g/L sodium bicarbonate, 10% fetal bovine serum (FBS), 50 IU/ml penicillin and 50mg/ml streptomycin. pT2A-BI-GapTdTomato-TRE-, pT2A-BI-GapTdTomato-TRE-SCF, or pT2A-BI-GapTdTomato-TRE-SDF-1 was transfected into Cos7 cells along with pT2K-M2-IRES2-NeoR^49^ and pCAGGS-T2TP using Lipofectamine 2000 according to the manufacture’s instruction. The cells were cultured G418-containing medium.

### *in vitro* under-agarose assay

This method was modified from the protocol reported by Heit *et al*.^21^. To control adhesion of PGCs with dish bottom, the surface of glass bottom dish (Eppendorf) was treated by 0.5 g/L of Pluronic®F-127 (Sigma-Aldrich) for 30 min. The treated dishes was dried by air for 15 min before use.

2 x Hank’s balanced salt solution (Gibco) and FAcs medium (1mM CaCl_2_) were mixed in 1:2 and warmed up to 70℃. UltraPure^TM^ Agarose (Invitrogen) solution (48 mg/ml) was prepared by microwave. HBSS/FAcs solution and UltraPure^TM^ Agarose solution were mixed in 1.99:1.7. 3 ml of mixed solution poured onto the glass bottom and solidified at room temperature (RT). 250 μl of FAcs was added to the gel and placed at the condition of 37℃, 5% CO_2_ for 30 min for equilibration. In case of chemical addition, BAPTA (Toronto Research Chemicals Inc), 2-APB (R&D Systems), SKF96365 (Wako), or ERseeing (Funakoshi) was added to the mixed solution of HBSS/FAcs and UltraPure^TM^ Agarose and each final concentration is 100 μM, 100 μM, 50 μM, and 0.3μM, respectively.

The gel was removed from the glass bottom dish. After removing the remaining solution on the dish, 2.5×10^5^ PGCs and 5.0×10^5^ FluoSpheres^TM^ Polystyrene Microspheres (Invitrogen, F8836) with a diameter of 10 μm were suspended in 5 μl of FAcs medium and seeded on the glass bottom dish and then the gel was put back on. 50 μl of FAcs and Doxycycline (Dox) were added on the gel. At this time, we adjusted the final concentration of Dox to 1 μg/ml.

### *in vitro* migration assay

1.4×10^4^ cells/μl of PGCs and 2.8×10^6^ cells/μl of Cos7 were suspended in 4-5% and 6-7% Matrigel® Basement Membrane Matrix Growth Factor Reduced, Phenol Red Free (Corning®)/FAcs (1mM CaCl_2_), respectively, and seeded next to each other of the glass bottom dish (Matsunami). The dish was warmed at 37℃ for 20 min to solidify Matrigel, and then FAcs (1mM CaCl_2_) and Dox (1 μg/ml at final concentration) were added on Matrigel. BAPTA, 2-APB, SKF96365, or ERseeing was added to FAcs and each final concentration is 100 μM, 100 μM, 50 μM, and 0.3μM, respectively.

### PGC-infusion into embryo

For back-infusion, cultured PGCs were collected by centrifugation at 100 g for 5 minutes with several washes in OPTI-MEM. The number of living PGCs was adjusted to 5,000 cells/μl, and 1 μl of such suspension was injected into the heart of HH15 chick or quail embryo by a fine glass capillary. For the tet-on induction, 1 μg/ml Doxycycline (Dox) (Clontech) was added into PGC-cultured FAcs medium 24 hours before the injection. For the Tet-on induction *in ovo,* a solution of Dox (0.5 ml of 100 μg/ml) was injected into the egg between the embryo and yolk^55^. They were incubated at 38.5℃ until intended developmental stage.

### Fixation, whole-mount and section immuno-staining

The embryos with infused PGCs were fixed with 4% PFA/PBS (paraformaldehyde/phosphate buffered saline) at 4℃ for 20 hrs. For immunostainings with QH1, EGFP, and mCherry in whole quail embryos, samples were washed with TNTT (0.1 M Tris-HCl (pH 7.5), 0.15 M NaCl, 0.05% Tween 20, 0.1% TritonX-100) (three times 30 min at RT). Blocking was performed in 1% blocking reagents (Roche)/TNTT for 1 hour at 4℃. The blocked samples were treated at 4℃ overnight with QH1 antibody (Hybridoma Bank, AB_531829, 1:200), anti-EGFP goat polyclonal antibody (GeneTex, GTX266673, 1:1,000), and anti-mRFP rabbit polyclonal antibody (ROCKLAND, 600-401-379, 1:1,000) in the blocking solution. After six 1-hour washes in TNTT at RT, the samples were treated with 1:500 anti-rabbit IgG-Alexa 555-conjugated donkey antibody (Invitrogen A31572), 1:500 anti-mouse IgG-Alexa 647-conjugated donkey antibody (Invitrogen A31571), and 1:500 anti-goat IgG-Alexa 488-conjugated donkey antibody (Invitrogen A11055) in 1% blocking solution at 4 ℃ overnight. Finally, they were washed six times for 1 hour each in TNTT at RT.

For section staining, fixed samples were washed with PBS twice, and treated with 30% Sucrose/PBS at 4℃ for 3-5 hrs. The samples were transferred into mixed solution of 30% Sucrose/PBS and O.C.T. Compound (Sakura Finetek Japan) (1:2) and treated at 4℃ overnight. After replacing the solution with O.C.T. Compound, the samples were freezed at -30℃. Frozen sections (12 µm thickness) were washed with TNTT three times, and treated with 1% blocking reagents /TNTT for 1 hour at RT. After blocking, the sections were incubated at 4℃ overnight using anti-EGFP goat polyclonal antibody (GeneTex, GTX266673, 1:1,000), and anti-mRFP rabbit polyclonal antibody (ROCKLAND, 600-401-379, 1:1,000) in the blocking solution. After three 5-min washes in TNTT at RT, the samples were treated with 1:500 anti-rabbit IgG-Alexa 555-conjugated donkey antibody (Invitrogen A31572) and 1:500 anti-goat IgG-Alexa 488-conjugated donkey antibody (Invitrogen A11055) in 1% blocking solution at RT for 1 hour, washed three times with TNTT, and mounted in Fluoromount-G^TM^, with DAPI (Invitrogen).

### Image acquisition and processing

For live imaging of *in vitro* under-agarose assay and *in vitro* migration assay, the samples on the glass bottom dishes (greiner bio-one) were placed on the incubation chamber (37°C, 5% CO_2_) (TOKAIHIT STX) equipped with the inverted microscope, IX83 (Olympus) interfaced to a spinning-disk confocal microscopy (Dragonfly200, OXFORD Instruments). Images were captured on a device camera and acquired using Fusion software (Dragonfly200, OXFORD Instruments). The acquired images were enlarged, edited and stacked in the Z-axis by the built-in function of Imaris (OXFORD Instruments).

For image acquisition of fixed whole-mount samples, images were obtained with the spinning-disk confocal microscopy (Dragonfly200, OXFORD Instruments). Images were captured on a device camera and acquired using Fusion software (Dragonfly200, OXFORD Instruments). Acquired Z-series images were deconvoluted and processed for 3D reconstruction by using Huygens (Scientific Volume Imaging) and Imaris software (ver.9.5, Oxford Instruments), respectively. These images were further reconstructed to 3D iso-surfaces with texture, and clipped with appropriate dorso-ventral plane(s).

For image acquisition of section samples, images were obtained with the inverted microscope, IX83 (Olympus).

### Quantification and statistical analysis

Areas and fluorescence intensities of GCaMP6s and mCherry in blebs and cell bodies in *in vitro* under-agarose assay were measured by ImageJ. Migration tracks and distances in *in vitro* migration assay were depicted and measured by the plug-in Manual trucking of imageJ. Fluorescence intensities of GCaMP6s and mCherry in front and rear areas in *in vitro* migration assay were measured by ImageJ. The number of protrusion-forming PGCs in *in vitro* migration assay were measured manually. For quantification of arrested PGCs in Ex-VaP 5 hrs after infusion, total area of EGFP or mCherry was measured by ImageJ. For quantification of PGCs in Ex-VaP 6 hrs after infusion and in the dorsal mesentery and gonads 2 days after infusion, the number of EGFP+ or mCherry+ PGCs was measured manually.

All box plots represent the mean, upper and lower interquartile, error bars (s.e.m) with median (x). p values were obtained by a 2-tailed, unpaired Student’s t test (Excel). All graphs were made by Excel.

## ACKNOWLEDGEMENTS

For 3D image processing, we thank the Center for Advanced Instrumental and Educational Support of the Faculty of Agriculture, Kyushu University. This work was supported by the following grants: JSPS KAKENHI (Grant number 22H02634 for D. S., and JP 19H04775 for Y. A.) and Princess Takamatsu Cancer Research Fund, Shinnihon Foundation of Advanced Medical Treatment Research, Terumo Life Science Foundation for D. S.

## AUTHOR CONTRIBUTIONS

M. Morita and D. S. designed the study. M. Morita performed most experiments and analyzed data. M. Morimoto performed the immunostaining experiments. T. T. performed 3D image acquisition. D. S., J. I. and Y. A. supervised the study. D. S. wrote the manuscript.

