## Supplemental Figures for "The SOCE system is critical for membrane bleb formation to drive avian primordial germ cell migration"

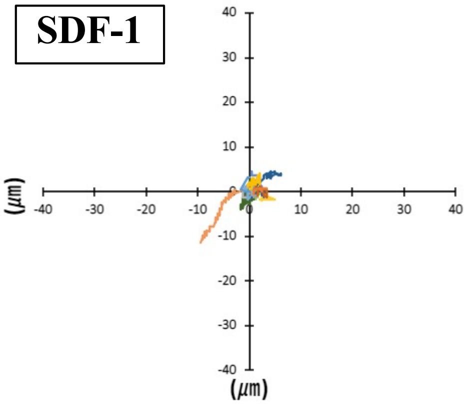


**Supplementary Figure 1 Morita *et al.***

**Figure S1 SDF-1 does not support PGC motility *in vitro***
Summary of PGC migration trajectories when co-culture with Cos7 co-expressing SDF-1 and Gap-TdTomato. Cos7 cells located in the left side. *N*=15 cells.

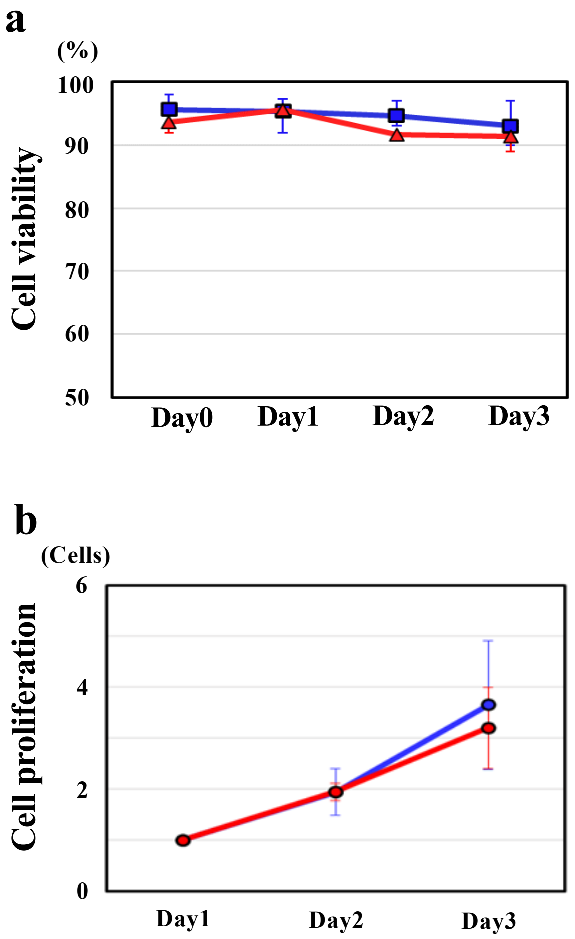


**Supplementary Figure 2 Morita *et al.***

**Figure S2 Orai1 E106Q+ PGCs show comparable survival and proliferation rates to control PGCs *in vitro***

**a** Comparison of cell viability rates between mCherry+ PGCs (control, blue line) and Orai1 E106Q+ PGCs (red line). *N*=3. **b** Comparison of cell proliferation between mCherry+ PGCs (control, blue line, *N*=26) and Orai1 E106Q+ PGCs (red line, *N*=29 cells). Dox is administrated to PGCs to induce expression of the dominant negative Orai1 at Day0. Error bars indicate standard error.
